# High-throughput isolation and culture of human gut bacteria with droplet microfluidics

**DOI:** 10.1101/630822

**Authors:** Max M Villa, Rachael J Bloom, Justin D Silverman, Heather K Durand, Sharon Jiang, Anchi Wu, Shuqiang Huang, Lingchong You, Lawrence A David

**Author notes:** denotes equal first-authorship (Max M Villa, Rachael J Bloom). **CORRESPONDING AUTHOR**: Lawrence A David, CIEMAS, Room 2171, 101 Science Drive, Box 3382, Durham, NC 27708.

## Abstract

Isolation and culture of gut bacteria enable testing for microbial roles in disease and may also lead to novel therapeutics. However, the diversity of human gut microbial communities (microbiota) impedes comprehensive experimental studies of individual bacterial taxa. Here, we combine advances in droplet microfluidics and high-throughput DNA sequencing to develop a platform for isolating and assaying microbiota members in picoliter droplets (MicDrop). MicDrop can be used to create millions of distinct bacterial colonies in a single experiment while using off-the-shelf parts compact enough to fit in an anaerobic chamber. In proof-of-concept experiments, we used the platform to characterize inter-individual metabolic variation among hundreds of polysaccharide-degrading gut bacteria from nine stool donors. We also used MicDrop to test the hypothesis that growth kinetics of individual gut bacterial taxa are associated with longterm community dynamics in an artificial gut. These demonstrations suggest the MicDrop platform could support future diagnostic efforts to personalize microbiota-directed therapies, as well as to provide comprehensive new insights into the ecology of human gut microbiota.

## Introduction

Bacterial culture was among the first techniques used to study human gut microbiota^1^. Bacterial isolation efforts beginning in the early 1900s identified key enteric genera such as *Bacteroides, Bifidobacterium*, and *Bacillus*^2^. Microbes isolated since then have served as crucial reagents for experiments. Gut bacterial isolates allow testing causal roles for specific microbes in animal models of metabolic and auto-immune disorders^3–5^. Bacterial isolates can also be genetically modified and tested *in vitro* to identify enzymatic machinery in processes like the fermentation of dietary fiber^6^, and cocktails of cultured bacteria are being explored as therapeutics for *C. difficile* infections and cancer^7–9^.

Yet a key challenge for current microbiota culturing efforts has been keeping pace with increasing knowledge and interest in gut microbial diversity. Culture-independent methods based on high-throughput 16S rRNA sequencing have revealed the average individual harbors hundreds of distinct enteric bacterial strains^10–13^. Moreover, unrelated individuals likely share no more than ~30% of bacterial strains^14^. Prevailing culture techniques do not scale to the diversity of microbes spanning human populations. Because most taxa are rare, exhaustive capture of bacterial species from even a single stool sample requires laborious spotting of thousands of bacterial colonies^15,16^. To reduce the human effort needed for such experiments, state-of-the-art culture assays leverage plate and liquid handling robots; but, even these automated efforts tend to be limited to tens of strains^17,18^. This limitation stems in part from the physical constraints of typical plate-based culture methods, which grow bacteria in wells ranging from centimeters to millimeters in diameter. Even relying on 96- and 384-well plates, conventional large-scale culture efforts may require loading and handling dozens of plates under anaerobic conditions^18^.

An alternative approach is to culture bacteria in small volumes (nano- to picoliters) by separating microbes into microscale wells. Devices composed of thousands of such wells have been used to culture both lab strains of bacteria and fungi^19^, as well as isolate previously uncultured bacteria from the gut and soil^20,21^. Even higher-throughput experiments are possible by compartmentalizing microbes in droplets of media that are tens to hundreds of microns in diameter and separated by immiscible oils and engineered surfactants^22,23^. Because droplets are not limited by the need to microfabricate physical wells or channels, millions of distinct culture volumes can be created on the order of minutes. Droplet techniques have so far been used to isolate uncultured microbes from seawater and soil communities^20,24,25^, assess microbial cross-feeding^26^, track population dynamics of individual bacteria^27^, and examine antibiotic sensitivity and commensal-pathogen interactions of human gut and oral microbiota^28,29^. Still, existing droplet microfluidic approaches for assaying bacteria have required combining complex emulsion techniques (water-oil-water) with flow cytometers or custom on-chip droplet sorting devices. These protocol requirements limit the accessibility of droplet technologies for bacterial assays and in their present form require equipment that does not fit into typical anaerobic chambers, which are needed to culture human gut bacteria^30^.

Here, we developed a platform to isolate and culture bacteria from human gut microbiota in droplets (MicDrop) using accessible techniques and equipment. A key challenge our method addresses is how to measure the growth of isolates within distinct microfluidic droplets. To accomplish this, we rely on 16S rRNA as intrinsic DNA barcodes that are shared between droplets carrying the same bacterial taxa, which we refer to here as a sequence variant or SV^31^. This approach in turn allows us to measure isolate growth in droplets without the need for double-emulsion techniques or droplet sorting. Instead, we combine single-emulsion (water-in-oil) microfluidic droplet protocols with molecular techniques (qPCR and 16S rRNA sequencing). These simplified protocols allow us to employ off-the-shelf microfluidic pumps and chips, which are compact enough to fit within typical anaerobic chambers. Using MicDrop, we characterized dietary polysaccharide metabolism among hundreds of gut bacteria from nine individuals. We then employed MicDrop to generate growth curves for dozens of distinct SVs in a single experiment, which we in turn used to investigate long-term microbiota dynamics of an artificial human gut. Together, these findings showcase the potential for microfluidic droplet techniques to characterize the growth and function of individual bacterial strains from complex gut microbial communities in high-throughput.

## Results

### MicDrop: a platform for culturing human gut microbiota in droplets

To isolate and culture individual gut bacteria from human gut microbiota, we merged concepts from prior microfluidic droplet protocols with high-throughput DNA sequencing (Fig. 1; Experimental Procedures). Our protocol first randomly encapsulates individual bacterial cells from gut microbiota into picoliter-sized droplets (Fig. 1A). Gut microbiota samples are diluted before encapsulation using the Poisson distribution at a loading concentration that optimizes the number of droplets loaded with cells (~10-26%) against the number of droplets loaded with more than one microbe (~95-86% of loaded droplets contain single cells) (Supplementary Fig. 1)^32^. Since many gut bacteria are obligate anaerobes, encapsulation takes place in an anaerobic chamber and droplets are subsequently incubated under anaerobic conditions (Supplementary Fig. 2). To track SV growth, we can avoid having to identify and sort bacteria by assuming that droplets are either empty or loaded with clonal isolates whose progeny share the same 16S ribosomal rRNA (rRNA) sequence, meaning genomic material accumulating across all droplets reflects the growth of SVs grown in isolation (Fig. 1A-D). We therefore track isolate growth in droplets at a given time point using bulk bacterial DNA extraction without droplet sorting, followed by DNA sequencing and total quantification (qPCR) of 16S rRNA. The product of relative SV levels from 16S rRNA sequencing and total 16S rRNA levels yields an estimate of the absolute levels of each SV across all droplets at the time of sampling.

**Figure 1.**
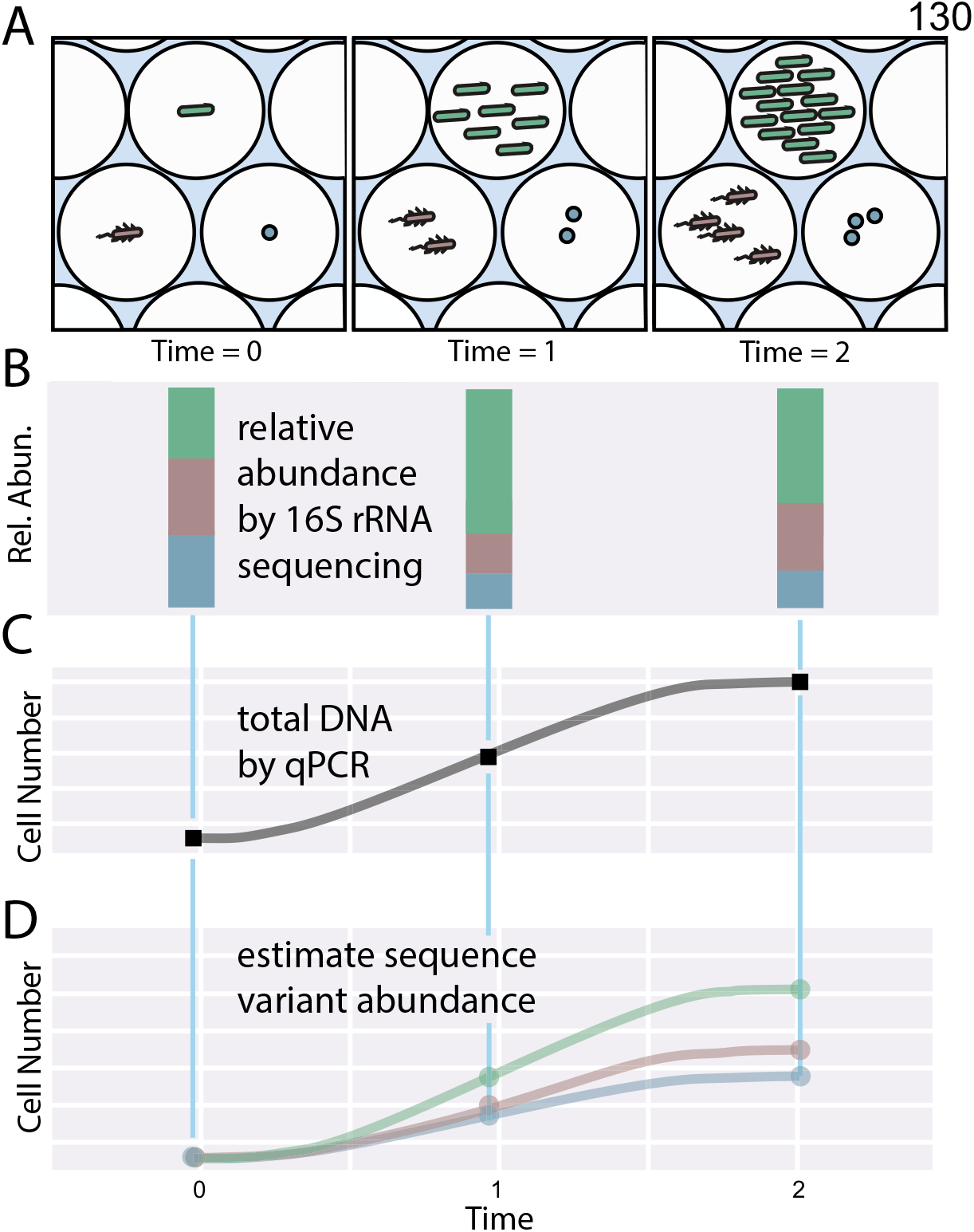
MicDrop concept. (A) Schematic of bacterial loading and growth in droplets over time. Bulk DNA is collected across all droplets and (B) the 16S rRNA gene is sequenced to establish the relative abundance of each sequence variant. (C) qPCR is used on the same samples to estimate the total abundance of all bacteria. Relative abundances are then multiplied by total bacterial levels to estimate sequence variant abundances (D).

To explore the feasibility of the MicDrop platform, we initially examined bacterial replication and isolation in droplets, as well as the use of DNA sequencing to track bacterial levels. We found aerobic monocultures of fluorescent *Escherichia coli* replicated in droplets in a qualitative manner that resembled growth on conventional Petri dishes (Supplementary Fig. 3). Droplet stability experiments suggested that bacteria could be studied in droplets for at least 5 days (Supplementary Fig. 4). Next, microscopy showed droplets could be used to segregate clonal isolate populations with distinct morphology and motility out of mixed microbial communities (Supplementary Fig. 5). Human fecal microbiota isolated and cultured in droplets exhibited 2.6 times more diversity than when grown in mixed conditions (Supplementary Fig. 6), which is consistent with the hypothesis that droplet isolation enables slow-growing microbes to be sheltered from competition with fast-growing bacteria^20^. Last, we found DNA-based techniques could be used to track bacterial levels in droplets. Quantitative growth measurements using qPCR of the 16S rRNA gene sampled every two hours from liquid cultures were also similar to *E. coli* grown on plates (ρ=0.95, p=8.7e-9; Spearman correlation; Supplementary Fig. 7). Next, using DNA sequencing, we analyzed a mixture of 10 bacterial strains that were isolated using MicDrop and grown under varying antibiotic conditions for 24 hours. We found that resulting bacterial DNA levels in droplets corresponded to isolates’ optical densities in reference well-plates (accuracy=75%; Supplementary Fig. 8).

### A droplet assay for prebiotic consumption by human gut microbes

To demonstrate how MicDrop could be applied to problems in human gut microbiology, we used the platform to measure bacterial utilization of carbohydrates. In typical carbohydrate utilization screens, bacteria are cultured in defined media containing a carbohydrate as the sole carbon source^17,33^. Microbes that replicate are assumed to be capable of utilizing the carbohydrate and are termed “primary degraders”^34,35^. The biology of primary degraders is of increasing interest because bacterial metabolism of select indigestible carbohydrates (prebiotics) leads to the growth and activity of gut microbes with multiple beneficial impacts on host health^17,36–40^. Still, bacterial prebiotic metabolism is incompletely understood, particularly with regards to the origins of wide inter-individual variation in microbiota metabolic potential^41–43^.

To explore how the MicDrop platform could assay bacterial prebiotic metabolism (Fig. 2A), we first loaded a previously characterized type strain *Bacteroides thetaiotaomicron* ATCC 29148 into microfluidic droplets and standard 96-well plates. Consistent with both prior studies^17^ and our well-plate experiments, *B. thetaiotaomicron* ATCC 29148 replicated in droplets on pullulan and levan, but not on lamanarin or a no carbohydrate control (Fig. 2B). We next tested how the MicDrop prebiotic assay performed using artificial microbial communities assembled from seven human gut isolates (Fig. 2E). Using 96-well plate experiments as our reference (Fig. 2C & 2D, Supplementary Fig. 9), we found the sensitivity, specificity, and false discovery rate of the MicDrop prebiotic assay to be 80%, 93%, and 9%, respectively (Supplementary Table 1). Finally, to assess the reproducibility of the MicDrop prebiotic assay, we used the same frozen fecal sample to compare the results of two separate experimental sessions. We observed higher correlation between replicates from the same session (*ρ*= 0.73-0.78, *p*<0.0001, Spearman correlation) than between replicates across sessions (*ρ*=0.57, *p*=1.67e-17, Spearman correlation). One explanation for the difference in correlation is that microbial communities re-assembled in different configurations each time microbiota was revived from frozen stool^44^. Indeed, controlling for microbiota differences between droplet inocula elevated the between session correlation (*ρ*=0.74, *p*=5.14e-19, Spearman correlation, Fig. 2F).

**Figure 2.**
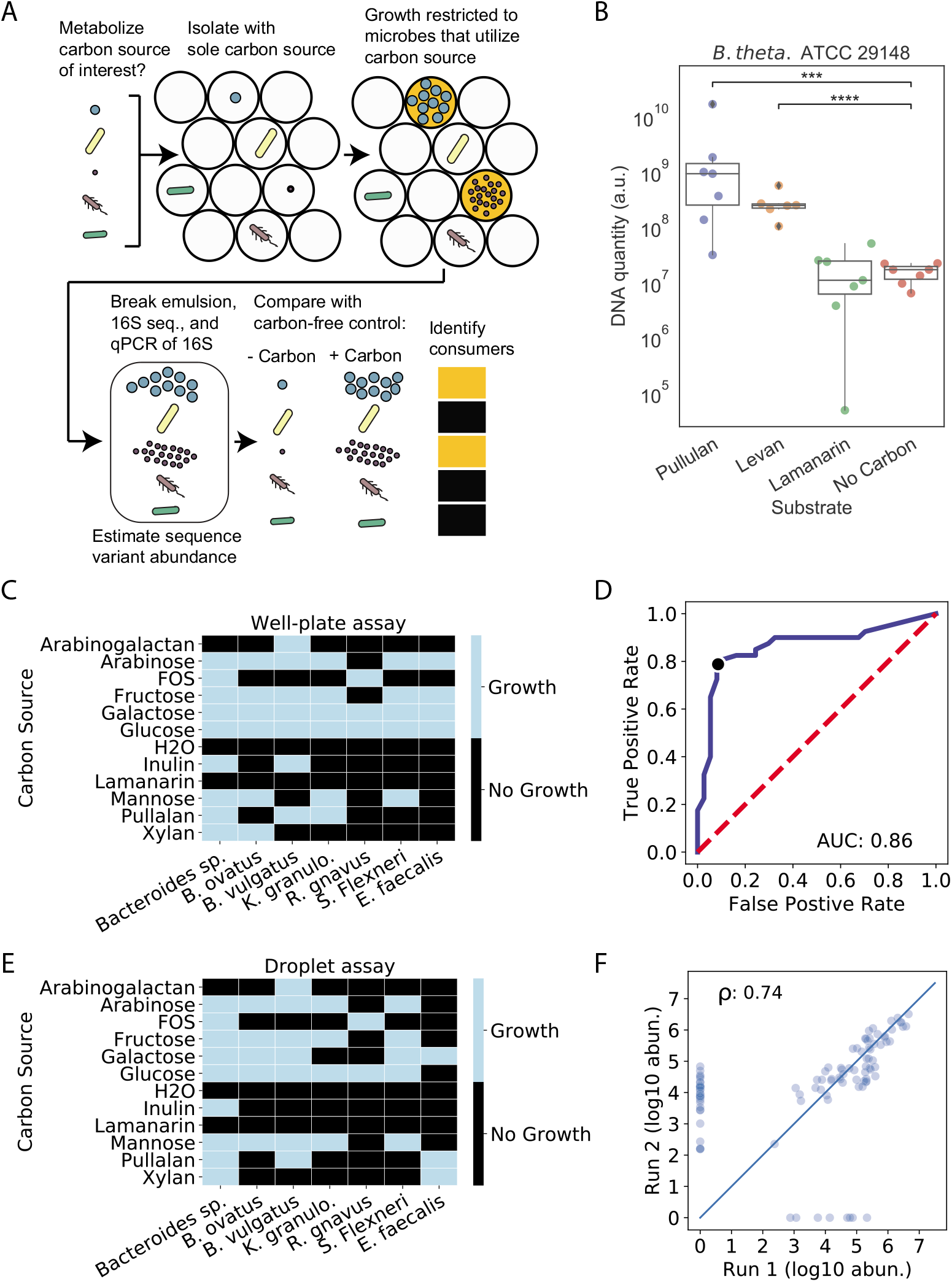
A prebiotic utilization screen based on the MicDrop platform. (A) Schematic of MicDrop prebiotic assay. (B) Droplet monoculture growth of *B. thetaiotaomicron* in microfluidic droplets measured by qPCR. (C) Results of 96-well plate growth of gut bacterial isolates across 11 carbohydrates. (D) ROC curve of MicDrop assay results at different growth threshold cut-offs using (C) as a reference. The black dot indicates the growth threshold that maximizes the true positive rate while minimizing the false positive rate, depicted in (E). (F) Correlation between two different MicDrop sessions (each carried out in triplicate) on the same frozen fecal sample and five different carbohydrates. Points indicate median growth of different SVs across each experimental session.

### Identifying primary degraders from human guts across multiple prebiotics

We next applied the MicDrop prebiotic assay to microbiota from nine healthy human stool donors. We assayed growth on three consumer-grade prebiotics (inulin, galacto-oligosaccharides (GOS), and dextrin) and a lab-grade prebiotic (xylan). Out of the 588 SVs detected across donor stool samples, MicDrop identified a total of 285 primary degraders that grew on at least one of the screened prebiotics (Fig. 3A). Prebiotic utilization patterns of primary degraders were similar to those of matched bacterial strains in a reference database of microbial carbohydrate utilization^45,46^. Among the instances when matched bacteria were annotated as consuming a prebiotic in the database, 86% were detected by MicDrop (*p*<0.001; permutation test). Still, we did find that 52% of instances when MicDrop indicated a primary degrader to consume a prebiotic were not reported for matched bacteria in the database (p=0.34; permutation test), which may in part reflect incomplete enumeration of bacterial carbohydrate utilization in the reference.

We next explored the hypothesis that differences in the presence or absence of primary degraders could drive inter-individual variation in human prebiotic response. Evidence arguing against this hypothesis included observing that multiple SVs capable of growing on the tested prebiotics were present in all subjects (median: 12.5 ± 6.1); Fig. 3B, Supplementary Table 2). Additionally, primary degraders were more likely to be shared between individuals than SVs not identified as primary degraders in stool samples (p=0.00122, Chi-square test). Concordant with human studies showing that even individuals with low prebiotic fermentation *in vivo* exhibit at least some prebiotic fermentative capacity *in vitro*^35^, our findings support the hypothesis that primary degraders are found across most individuals.

Still, we also found evidence for inter-individual variation in primary degrader composition and abundance. We observed differences in the richness of primary degraders across subjects (*p*<0.001, Two-way ANOVA; Fig. 3B). We also found subject identity explained more variation (R^2^=0.30, PERMANOVA; Supplementary Table 3) than prebiotic type (R^2^=0.16, PERMANOVA; Supplementary Table 3) in overall primary degrader growth. Last, we observed differences in the relative abundance of primary degraders in inoculating fecal communities (*p*<0.0001, Two-way ANOVA; Fig. 3C).

Thus, while primary degraders are likely present in most individuals, differences in polysaccharide metabolism may be due to inter-individual variation in primary degrader abundance in the gut^41,42^.

**Figure 3.**
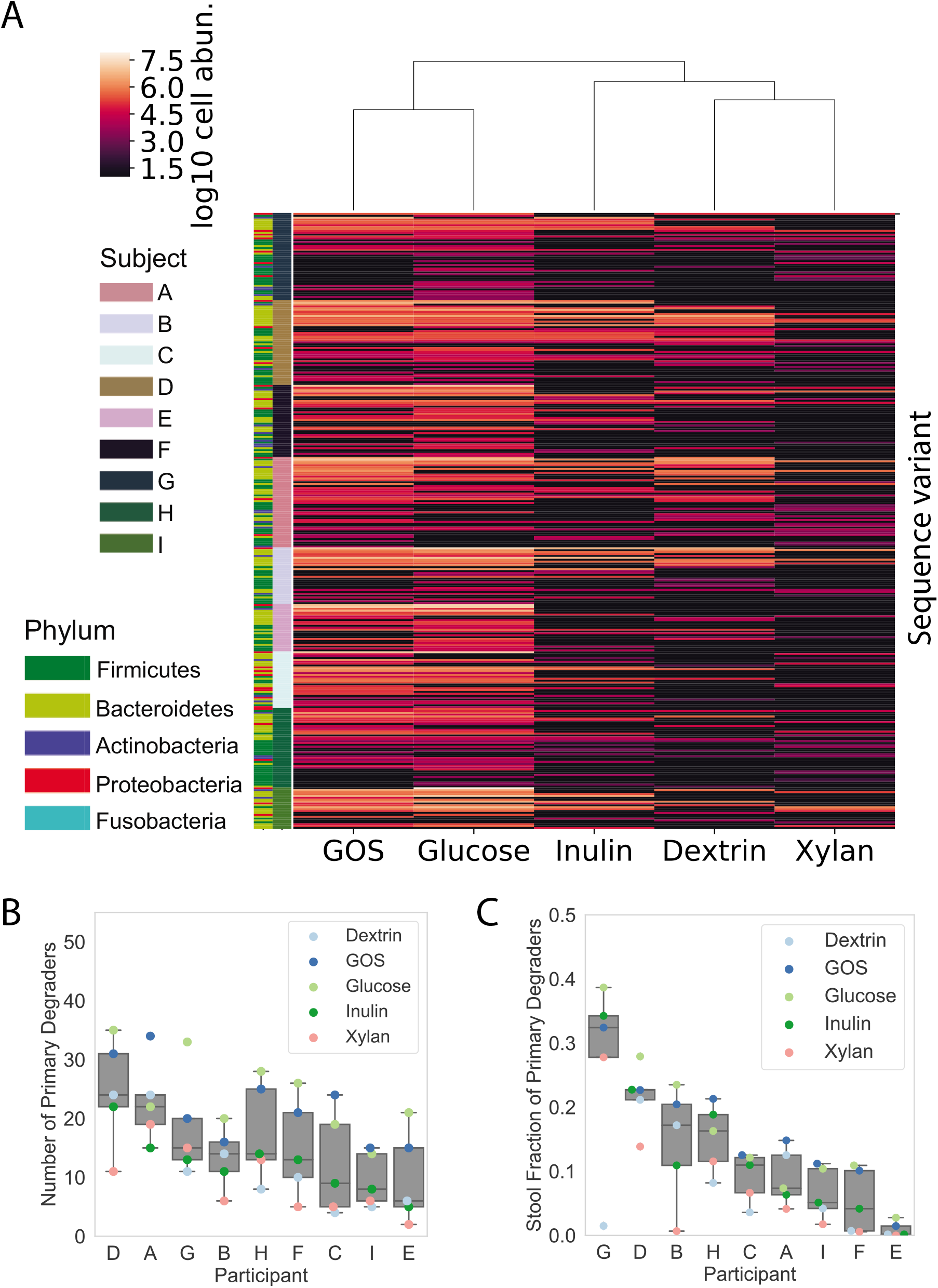
MicDrop prebiotic assay carried out on fecal samples from nine individuals. (A) Microbial carbohydrate preferences for 328 SVs from nine healthy human donors. Of these SVs, 285 grew on at least one of the prebiotics (*i.e.* GOS,inulin, dextrin, or xylan). (B) The number of primary degraders detected by MicDrop and (C) the relative abundance of primary degraders in stool samples differed by participant and prebiotic (*p*<0.001, Two-way ANOVA). Participant orderings in (B) and (C) are sorted by median values.

### High-throughput estimation of growth dynamics of human gut microbiota

In a second set of demonstration experiments, we used the MicDrop platform to explore the dynamics of human gut microbiota in artificial gut systems. Such systems have been used when *in vivo* microbiota research is challenging, including measuring the effects of nutrition on the infant gut, systematic antibiotic testing, and investigating chemotherapy-induced dysbioses^47,48,49^. Still, a recurring challenge of artificial gut models has been their inability to completely reconstruct *in vivo* microbial communities. After inoculation, these models often decrease in diversity within 24 hours^50–52^, and as little as 15% of the starting community may ultimately remain after a week of culture^53^. A potential explanation for part of this diversity loss is that bacteria are sensitive to media conditions: studies using individual gut bacterial strains show growth can be affected by even a single medium component^54^; and, varying media used in artificial guts leads to broad scale changes in microbial community structure^55,56^. Still, the hypothesis that individual gut microbes’ suitability to media is associated with their long-term persistence in artificial gut models has not been fully explored, likely due in part to the challenges of isolating and assaying each component species in these communities.

To test the hypothesis that growth of individual microbial SVs in a particular medium would correspond to SV persistence in an artificial gut setting, we used stool from a healthy human donor to inoculate a continuous flow bioreactor system we have used in past gut microbial ecology studies^57,58^. The artificial gut was supplied with modified Gifu Anaerobic Medium (mGAM)^59^, which features a variety of carbon and nitrogen sources, as well as extra amino acids, vitamin K, and hemin. We chose mGAM because it enables a wide growth of mammalian gut bacteria^54,59^. Yet, despite this choice of medium, microbiota dynamics in the artificial gut exhibited the same loss in diversity observed in prior studies (Supplementary Fig. 10)^50,53^. At the end of two weeks of culture, only 23% and 18% of inoculating bacteria genera and SVs, respectively, were still detected in the artificial gut (Supplementary Table 4).

A fresh stool sample from the same donor used to inoculate the artificial gut was then assayed by the MicDrop platform. To additionally demonstrate the potential for MicDrop to assess the kinetics of bacterial growth over time, we created replicate droplet populations and destructively sampled them at hourly intervals for the first 24 hours, and daily for four subsequent days after inoculation. Among the resulting time series, 94 SVs were detectable in droplets, meaning they appeared in >5 time points; Table 1, Supplementary Fig. 11). These SVs included representatives from the major human gut bacterial phyla (the Actinobacteria, Bacteroidetes, Firmicutes, and Proteobacteria) and represented 76% of the inoculum’s SVs, a proportion approaching prior culture efforts using mGAM medium^59^. Of the detectable SVs, we then measured how many exhibited evidence for growth in droplets. We defined a cut-off for growth as inferred doublings of at least 2.14 times (Δ ln(SV DNA abundance) ≥ 1.48) based on our antibiotic-based control experiments (Supplementary Fig. 8). A total of 34 SVs were defined as growing (Fig. 4A), which accounted for 25% of the inoculum’s SVs. Of the SVs with positive growth, 12 SVs were not detected by sequencing in the inoculum, which suggests they could be laboratory contaminants. Still, these SVs resemble known gut bacteria (Supplementary Table 5) and may alternatively represent rare microbes that require culture to be detected, which is a previously reported phenomenon^60^.

**Table 1.** Number and fraction of microbes from a human stool sample cultured by MicDrop in mGAM medium. SVs were considered as ‘detected’ if present in more than five longitudinal measurement. ‘Growth’ was defined by an inferred number of doublings equal or greater than 2.14.

| Taxonomic level | Taxa in inoculum | Taxa detected in droplets | Taxa in inoculum & detected in droplets | Fraction of inoculum detected in droplets | Taxa grew in droplets | Taxa in inoculum & grew in droplets | Fraction of inoculum that grew in droplets |
| --- | --- | --- | --- | --- | --- | --- | --- |
| **Phylum** | 4 | 5 | 4 | 1.00 | 5 | 4 | 1.00 |
| **Class** | 10 | 11 | 10 | 1.00 | 7 | 5 | 0.50 |
| **Order** | 10 | 14 | 10 | 1.00 | 8 | 5 | 0.50 |
| **Family** | 17 | 21 | 16 | 0.94 | 13 | 7 | 0.41 |
| **Genus** | 56 | 53 | 40 | 0.71 | 20 | 13 | 0.23 |
| **Sequence Variant** | 89 | 94 | 68 | 0.76 | 34 | 22 | 0.25 |

**Figure 4.**
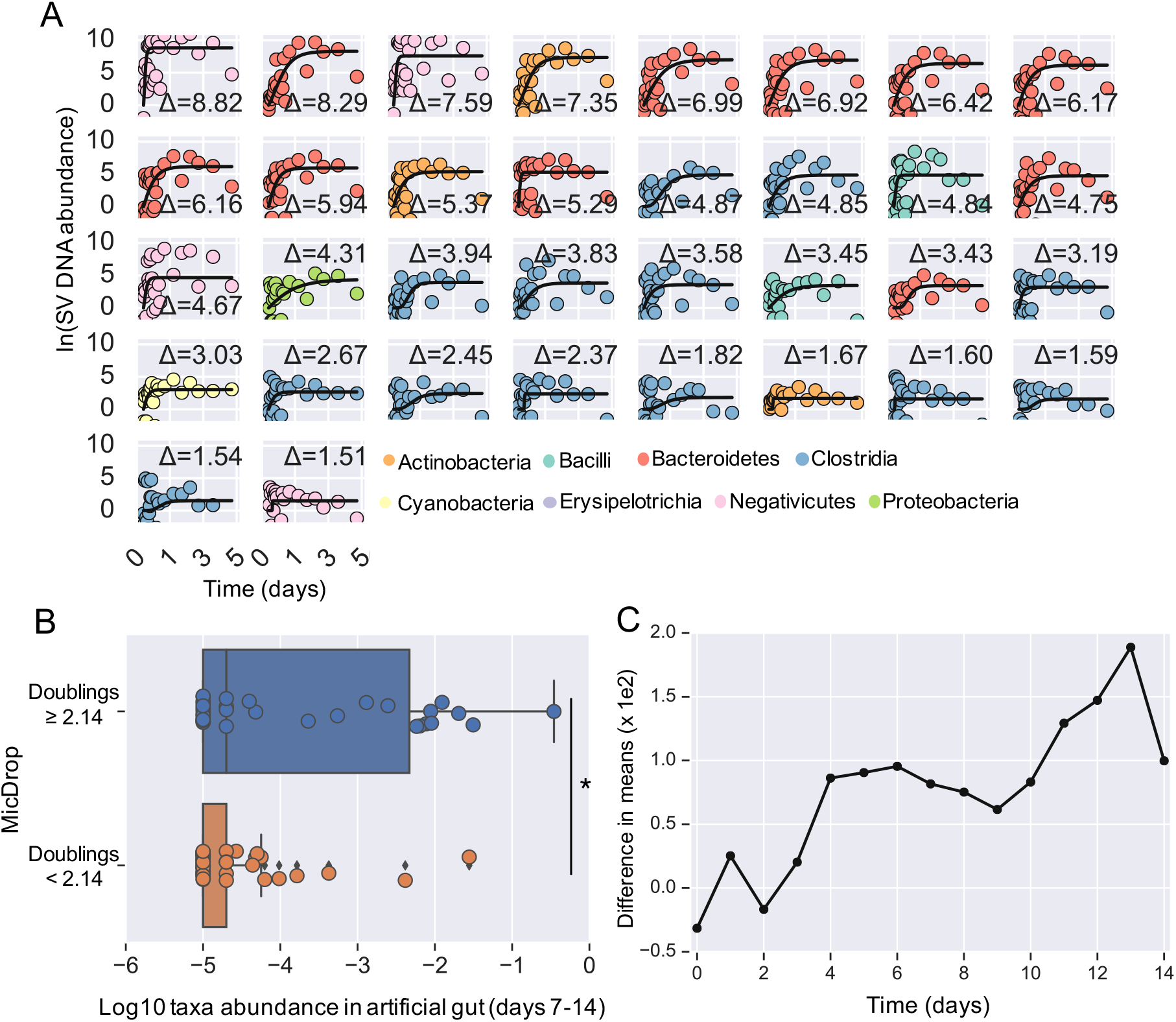
Comparison of SV growth kinetics and persistence in an artificial gut. (A) Abundance over time of SVs in MicDrop from a fresh human fecal sample. Modified Gompertz growth curves are fit to time-series. SVs are colored by taxonomy and sorted according to total growth (curve asymptote height; indicated by Δ), which is denoted on each sub-plot. Only SVs inferred to double at least 2.14 times were considering growing and are shown (ln(Δ SV DNA abundance) ≥1.48; threshold determined using control experiments in Supplementary Fig. 8). To ease viewing, curves are shifted vertically so y-intercepts are at the origin. (B) Long-term abundance of SVs in an artificial gut (grouped across days 7-14) grouped by whether SVs were identified by MicDrop as growing (doubling ≥ 2.14 times) or non-growing doubling < 2.14 times) (Mann-Whitney U, *p* < 0.02). (C) Differences in mean abundances of growing and non-growing SVs increased over time in an artificial gut system (*ρ*=0.80, *p* < 0.0004, Spearman correlation).

The growth of SVs measured with MicDrop were ultimately associated with SV dynamics in the artificial gut. Such associations were not apparent on short-time scales (i.e. 1-5 days after inoculation), which is consistent with the notion that non-growing SVs require several days to wash out of an artificial gut after inoculation. However, from day 7-14 of the artificial gut experiment, we observed elevated abundances among artificial gut SVs that grew in the MicDrop platform (inferred doublings ≥ 2.14) relative to ones that did not (SV doublings < 2.14) (*p* < 0.02, Mann-Whitney U test) (Fig. 4B). This difference in abundance increased over time in the artificial gut system (*ρ* = 0.80, *p* < 1e-4, Spearman correlation) (Fig. 4C). Still, some SVs grew well in droplets, but did not persist in the artificial gut (left-most points of upper bar in Fig. 4B); or, by contrast, did not grow in droplets, but were relatively abundant in the artificial gut (right-most points of lower bar in Fig. 4B). The former may represent examples of SVs that are outcompeted in mixed culture, while the latter may be examples of SVs that depend on inter-species interactions to persist.

## Discussion

We report here a microfluidic platform for isolating, culturing, and assaying component members of human gut microbiota (MicDrop) using accessible microfluidic and molecular techniques. We used MicDrop to compare the growth kinetics of dozens of microbial SVs to the dynamics of an artificial gut community and to examine interindividual variation in gut bacterial polysaccharide metabolism. The flexibility of the platform suggests its underlying concepts could be applied to assaying microbial responses to other compounds including pharmaceuticals, antibiotics, or host-secreted compounds^18,61^ using individual members of communities comprised of microbes from culture collections, mutant libraries, other human body sites, or environmental systems. The ability of MicDrop to screen clonal populations could be particularly useful for assays characterizing the behavior of isolates free from the effects of inter-species interactions like competition or facilitation^17,18^.

Yet, we acknowledge MicDrop still has some limitations. We rely on 16S rRNA as a molecular barcode for droplets sharing the same bacterial SV, meaning that the platform is sensitive to similar challenges due to inter-species rRNA copy number variation confronting 16S rRNA microbiota surveys^62^; and MicDrop cannot detect differences in growth originating from distinct clones of the same SV. For precise growth assays targeting bacteria from a limited number of taxa, traditional culture methods could be better suited. An additional limitation of MicDrop in its current form is the time and manual effort needed to setup individual droplet generation experiments and ensure accurate Poisson dilution of bacterial cells. Experimental effort could be reduced and reproducibility enhanced by automating sample switching. Last, we focused here on culture in liquid media using soluble substrates; future extensions of MicDrop that provide solid physical surfaces to colonize^63,64^ or insoluble substrates like mucin will require developing techniques to avoid the clogging of microfluidic channels.

Still, in its present form, MicDrop enabled useful insights into human gut microbiology. Our findings that bacterial SV growth in isolation is associated with persistence in an artificial gut supports the ecological hypotheses that intrinsic lifestyle characteristics of bacteria shape overall community dynamics. Indeed, species’ growth rate (measured by 16S rRNA copy number) is positively correlated with microbes’ relative abundance in seawater^65^, as well as skin microbiota of amphibians^66^. Additionally, our microfluidic investigation of prebiotic response supports hypotheses that inter-individual variation to carbohydrate interventions is due to differential abundances of polysaccharide degrading bacteria between people^43,67^. Droplet microfluidics could be used in the future to stratify human populations into groups most likely to benefit from prebiotic treatments^41^, by providing a culture-based diagnostic approach capable of scaling to the diversity of microbes inhabiting the human gut.

## EXPERIMENTAL PROCEDURES

### Overall MicDrop procedure

Droplets were made on a microfluidic chip (6-junction droplet chip, Dolomite

Microfluidics). Bacterial media varied by assay; for the oil phase, we used a fluorinated oil and surfactant mixture 1% Picosurf (Sphere Fluidics) in Novec 7500 (3M). One day prior to performing the droplet assay, all reagents including carrier oil, culture media, and carbon solutions were equilibrated to the anaerobic atmosphere in an anaerobic chamber (Coy). The fecal inoculum optical density at 600 nm was recorded and diluted according to the Poisson distribution: 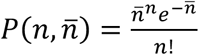, where *n* is the droplet occupancy (i.e. 0,1,.. cells/droplet) and 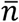 is the average number of cells per droplet given by: 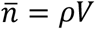 where V is droplet volume and *ρ* is cell density. Assays were performed at a 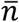 of 0.1-0.3 to minimize the number of droplets loaded with more than one cell (Supplementary Fig. 1). Thus, for a fixed droplet volume and 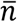, the target cell concentration can be obtained from: 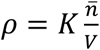, where *K* is a constant that converts CFUs/mL to OD_600_ determined from replicate CFU assays. Syringe pumps were used to control the flow rates of the oil and cell suspension (NE-1000 Single Syringe Pump, New Era Pump Systems). Following the culture period, droplets were loaded into chambered slides (C10283, Invitrogen) or directly onto glass slides and observed with Phase and/or Darkfield microscopy (Nikon) to examine growth and the appropriate loading. All steps of cell encapsulation and culture were performed in an anaerobic chamber.

### Collection and preparation of fecal inoculum for artificial gut and MicDrop growth dynamics assays

Stool was collected from human donors under a protocol approved by the Duke Health Institutional Review Board (Duke Health IRB Pro00049498). Inclusion criteria limited this study to healthy subjects who could provide fecal samples at no risk to themselves, had no acute enteric illness (e.g. diarrhea) and had not taken antibiotics in the past month. Fresh stool samples were collected in a disposable commode specimen container (Fisher Scientific, Hampton NH). Intact stool was moved within roughly 15 minutes of bowel movement into anaerobic conditions. The sample was prepared for inoculation in an anaerobic chamber (Coy). A 5 g stool aliquot was weighed into a 7 oz filtration bag (Nasco Whirl-Pak) and combined with 50 mL of mGAM media (Gifu Anaerobic Medium, HiMedia, with the addition of 5 mg/L Vitamin K and 10 mg/L Hemin,^59^) that was pre-reduced overnight in an anaerobic chamber. The mixture was homogenized in a stomacher (Seward Stomacher 80) on normal speed for 1 minute under atmospheric conditions to make a total of 100 mL of inoculum. The supernatant was decanted into beakers and loaded into syringes for inoculation into the artificial gut or filtered through a 50 μm filter (Celltrics) and diluted and loaded into droplets.

### Droplet DNA extraction, PCR amplification, and DNA sequencing

To extract DNA from droplets, excess oil was removed by pipetting and water-in-oil emulsions were broken by adding an equal amount of 1H,1H,2H,2H-Perfluoro-1-octanol (PFO, VWR) and briefly vortexed. Then, the samples were briefly centrifuged (<200 g) to separate the aqueous and oil phases by density. The aqueous solution was transferred to a new tube, and DNA was extracted using a kit (Qiagen #12224). DNA was extracted from artificial gut and stool samples using a 96-well PowerSoil kit (Qiagen #12888). For all samples, the V4 region of the 16S rRNA gene was barcoded and amplified from extracted DNA using with custom barcoded primers, using published protocols^68,69^. 16S rRNA amplicon sequencing was performed on an Illumina MiniSeq with paired-end 150 bp reads. We chose to only analyze samples with more than 5,000 reads to remove outlying samples that may have been subject to library preparation or sequencing artifacts. The 16S rRNA nucleotide sequences generated in this study will be made available at the European Nucleotide Archive under study accession number TBD. Total bacterial abundances from droplet cultures were estimated by qPCR for bacterial 16S rRNA using the same primers used in the DNA sequencing protocol. Amplification during the qPCR process was measured with a Real-Time PCR system (CFX96 Real-Time System, BioRad) using *E. coli* DNA at a known cell concentration as a reference.

### Identifying Sequence Variants and Taxonomy assignment

DADA2 was used to identify SVs^31^. Custom scripts were used to prepare data for denoising with DADA2 as previously described^57^. Reads were then demultiplexed using scripts in Qiime v1.9^70^. SVs were inferred by DADA2 using error profiles learned from a random subset of 40 samples from each sequencing run. Bimeras were removed using the function removeBimeraDenovo with tableMethod set to “consensus”. Taxonomy was assigned to sequence variants using a Naïve Bayes classifier^71^ trained using version 123 of the SILVA database^72^. For growth dynamics of the human gut microbiota and microbiota dynamics in the artificial gut, only forward sequencing reads were analyzed. Downstream analysis on sequence variant tables was performed using R (ver. 3.4.2) and Python (ver. 2.7.6). PERMANOVA was run in R using adonis in the vegan package (ver. 2.5-2).

### Growth dynamics of human gut microbiota

To estimate SV growth curves using MicDrop, we collected a total of 70 separate microfluidic droplet aliquots for destructive longitudinal sampling. Droplets were generated according to the MicDrop protocol described above. We used a modified Gifu Anaerobic Medium (mGAM) in our droplets (Gifu Anaerobic Medium, HiMedia, with the addition of 5 mg/L Vitamin K and 10 mg/L Hemin). Each aliquot of 200 μl of droplets was incubated at 37 °C in an anaerobic chamber. Aliquots were destructively sampled in triplicate, hourly, for hours 0-24 after droplet making and in duplicate once a day for hours 24-127 after droplet making.

Growth curves were fit using a combination of 16S rRNA qPCR and DNA sequencing data. To minimize the potential for poorly fit growth curves, SVs were required to have been detected by DNA sequencing in >5 samples to be included in curve fitting. To avoid numerical instabilities associated with taking the log or dividing by zero, a pseudocount of one was added to the sequence variant count table prior to normalization to relative abundances. Relative abundances of each SV were then determined by dividing the number of counts associated with each SV in each sample by the total read counts in the sample. Concentrations of each taxa were then estimated by multiplying the relative abundances of SVs by the 16S rRNA concentrations determined by qPCR. Technical replicates constituted distinct data points in these calculations. We used the SciPy Python package (v0.19.1) to fit a modified Gompertz equation^73^ to which we added an additional term to account for differences in starting abundance to the resulting dataset: 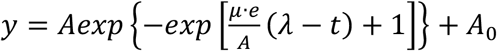, where μ is growth rate, A is carrying capacity, λ is lag time, or the time it takes for a bacteria to reach logarithmic growth, and A_0_ accounts for the relative abundance of different SVs in the inoculum. We fit curves using the module scipy.optimize.least_squares with the robust loss function “soft_l1”. Parameter bounds were also used to minimize the optimization search space. We set lower bounds of A=0, λ=-50, μ=0, A_0_=0; and, upper bounds of A=15, λ=12, μ=2.6, A_0_=15. We selected bounds by considering both biological feasibility and parameter sensitivity analyses (Supplementary Fig. 12). Our upper bound for growth rate (μ=2.6) represented a doubling time of 15 minutes, which we based on the fastest growth rates observed in an anaerobic bacterium^74^. The upper bounds on *A* and *A*_0_ were set to the maximum amount of DNA measured across replicate MicDrop samples from the human fecal inoculum. The upper bound on λ, which represents the lag time until exponential growth^75^ was set at 12. Lower bounds of 0 for *A, A0*, and μ reflect our choice not to model negative growth. A lower bound for λ was selected by sensitivity analysis (Supplementary Fig. 12), which revealed that a bound of zero led to fitted λ values regularly collapsing to our boundary limits. We also found that fitted curves were sensitive to starting parameters. To ensure a broad search of parameter space, we initialized each curve fit multiple times (n=100) with starting parameters randomly distributed between the bounds of each parameter. Fitted growth rates often collapsed to the maximum μ tolerated; we therefore only retained fits where growth rates were at least slightly below our upper bound for μ (μ < 2.5) (Supplementary Fig. 13). Of the remaining fitted curves, we analyzed the one with the lowest loss function. In our analyses of SV growth in human fecal samples, we defined total SV levels as *y*(127 hours) – *y*(0 hours).

### Microbiota dynamics in an artificial human gut

We cultured human gut microbiota with an artificial gut model that we have used in prior studies^57,76^. A continuous-flow artificial gut system (Multifors 2, Infors) was used to culture gut microbiota seeded from human stool. A vessel was autoclave-sterilized and prepared with 300 mL of mGAM media (see *Growth dynamics of human gut microbiota* above). We inoculated the vessel with 100 mL of fecal inoculum, resulting in a total culture volume of 400 mL. After 24 hours, the media feed was initiated at a constant rate of 400 mL per day. A carboy feeding the media was changed once over the course of the 14 days. Feed rate, oxygen, pH, temperature, and stir rate were all controlled by software (IRIS v6, Infors). Positive pressure within the vessels was maintained to prevent contamination by sparging with nitrogen at 1 LPM. Dissolved oxygen concentration was measured continuously using Hamilton VisiFerm DO Arc 225 probes. pH was monitored with Hamilton EasyFerm Plus PH ARC 225 probes and was maintained between 6.9 and 7.1 using a 1 N HCl solution and a 1 N H_3_PO_4_ solution.

The vessel was maintained at 37°C via the Infors’ onboard temperature control system. The vessel was continuously stirred at 100 rpm using magnetic impeller stir-shafts. Samples were taken once every 24 hours between 1 PM and 5 PM for 14 days and were frozen immediately at −80°C for later extraction.

### MicDrop Prebiotic Assay

Microbial communities isolated from human stool samples were tested for carbohydrate consumption using the MicDrop platform. Cells were revived from frozen stock in rich medium (mGAM, see *Growth dynamics of human gut microbiota)* for 18 hours to allow cells to recover from freezing. Bacteria were then cultured in minimal medium (Supplementary Table 6) containing glucose and galactose (Sigma) as the sole carbon sources to deplete excess nutrients^77^. Following determination of the loading concentration, the bacteria were washed twice by centrifugation (2 min at 14,000 g) to remove free monosaccharides and resuspended in 2X minimal medium without a carbon source. Bacteria were filtered using a 50 μm filter (CellTrics, Sysmex) to remove multi-cell clumps. The filtered microbiota suspension was then added to prebiotics in a 50:50 mixture of 1% prebiotic solution and 2X minimal medium. To prevent chip fouling during droplet generation, the oil inlet was equipped with 10 μm inline filters (P-276, IDEX). Droplet generation in the anaerobic chamber was monitored using a bright field microscope (Celestron). Droplet cultures were stored in 5 mL polypropylene tubes (Falcon) with the caps closed in an anaerobic incubator at 37 °C. Following the second day of incubation, cultures were moved to a −20 °C freezer for storage prior to DNA extraction.

### Validation of prebiotic utilization assays

To validate the MicDrop prebiotic assay, we generated reference data on carbohydrate preferences using an artificial community of seven wild-type gut isolates from our culture collection (Supplementary Table 7), which were grown in both 96-well plates and the MicDrop Prebiotic Assay described in the preceding paragraph. Following the same procedure described in the *MicDrop Prebiotic Assay*, well plates were prepared with minimal medium (Supplementary Table 6) and a carbohydrate as a sole carbon source (Supplementary Table 8). A 10 μL aliquot of bacterial suspension was added to 200 μL of medium in 96-well plates and incubated in a humidified container for two days at 37°C. All culture experiments were performed in an anaerobic chamber. Following the culture period, the optical density at 600 nm of each well was examined using a plate reader (CLARIOstar, BMG Labtech). Following published protocols^17^, isolate growth in plates was normalized to the maximum growth for each microbe. To classify isolates as either “growing” or “not-growing,” a threshold of 20% of maximum growth was applied to the plate data, above which was considered growth on the carbon source of interest. The same isolates used in the well-plate analyses were mixed evenly into an artificial community and examined using the MicDrop prebiotic assay described above. MicDrop experiments were performed in triplicate. Growth thresholds for the MicDrop assay were determined by first pre-processing sample qPCR values to zero if they indicated overall growth below mean no-carbon controls. Then, relative SV abundance data were converted to absolute SV abundances by multiplying each sample by the corresponding qPCR value. Median SV abundances were then calculated across replicates and SV abundances from matched no-carbon controls were subtracted from each sample. An optimal SV growth threshold for determining growth on a carbohydrate in MicDrop was determined by applying Youden’s J index across all possible threshold values, with the well-plate data as the reference (Supplementary Fig. 9). A growth threshold of 88% maximized this index and was used in subsequent experiments on fecal samples.

### MicDrop prebiotic assays using human stool samples

Stool samples were collected from nine healthy donors (7 men, 2 women) between the ages of 35-53 under the IRB protocol described in section *Collection and preparation of fecal inoculum for artificial gut and MicDrop growth dynamics assays.* To facilitate carrying out prebiotic assays simultaneously across a range of donors, we used frozen gut microbiota in these experiments. Fecal slurries were made at 10% w/v using mGAM medium and a stomacher (Seward) that homogenized fecal samples for one minute. Then, slurries were mixed 50:50 with 50% glycerol and stored at −80 °C for later use. Samples were assayed following the MicDrop prebiotic assay procedure described above.

### Comparison of primary degraders to the Virtual Metabolic Human database

We compared the identity of primary degraders to carbon consumption profiles from the Virtual Metabolic Human (VMH) database^46,78^. We first mapped primary degraders to this database by taking SV 16S rRNA sequences from our study and searching for matches in the NCBI nucleotide database. Each 100% match was then linked to a type strain in the VMH database using NCBI taxonomy ids. Since some NCBI taxonomy IDs could be mapped to multiple strain IDs, we used an NCBI genome assembly file (ftp://ftp.ncbi.nlm.nih.gov/genomes/ASSEMBLY_REPORTS/assembly_summary_genbank.txt) to perform more specific mappings; when more than one mapping was possible, we selected the strain with the oldest genome annotation (we reasoned that strains that were selected first for sequencing were also more likely to have more thorough experimental characterizations). Once SVs were mapped, we restricted our analysis to SVs that grew on a given carbohydrate in over half of the participants and where all BLAST matches to the VMH database had concordant prebiotic utilization annotations. Only consumption of xylan and inulin were examined since GOS and dextrin were not referenced in the VMH database. Permutation analysis were carried out by randomly shuffling rows and columns of the droplet primary degrader table and repeating the analyses.

## Acknowledgements

The authors would like to thank Christopher Mancuso and Ahmad S. Khalil, Ph.D., for their helpful comments on the manuscript. L.A.D. acknowledges support from the Global Probiotics Council, a Searle Scholars Award, an Alfred P. Sloan Research Fellowship, the Beckman Young Investigator program, the Translational Research Institute through Cooperative Agreement NNX16AO69A, the Damon Runyon Cancer Research Foundation, the UNC CGIBD (NIDDK P30DK034987), and NIH 1R01DK116187-01. This work used a high-performance computing facility partially supported by grant 2016-IDG-1013 (“HARDAC+: Reproducible HPC for Next-generation Genomics") from the North Carolina Biotechnology Center. *E. coli* strain was provided by N. Lord and J. Paulsson. M.M.V. holds a Postdoctoral Enrichment Program Award from the Burroughs Wellcome Fund. This material is based upon work supported by the National Science Foundation Graduate Research Fellowship under Grant No. DGE-1644868 to R.J.B.

## Author contributions

M.M.V., R.J.B., and L.A.D. developed the assay platform and designed the study. M.M.V., R.J.B., H.K.D, S.H., S.J. and A.W. performed the experiments. J.D.S. contributed software for bioinformatic analysis. M.M.V., R.J.B., and L.A.D. analyzed the data. S.H. and L.Y. helped with design and data interpretation. M.M.V., R.J.B., and L.A.D. wrote the manuscript with feedback from all authors.

