## Supplementary_information for "High-throughput isolation and culture of human gut bacteria with droplet microfluidics"

**SUPPLEMENTAL FIGURES**

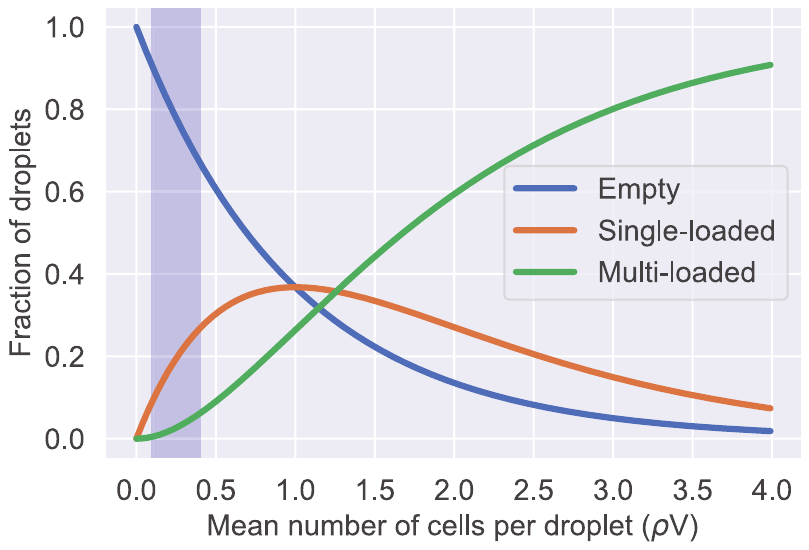
 **Supplemental Figure 1. Poisson loading distributions.** Tradeoffs exist between the number of droplets loaded with bacteria and how many of those loaded droplets are clonal. MicDrop protocols here balance these tradeoffs by loading droplets at means ranging from 0.1-0.3 (highlighted region).

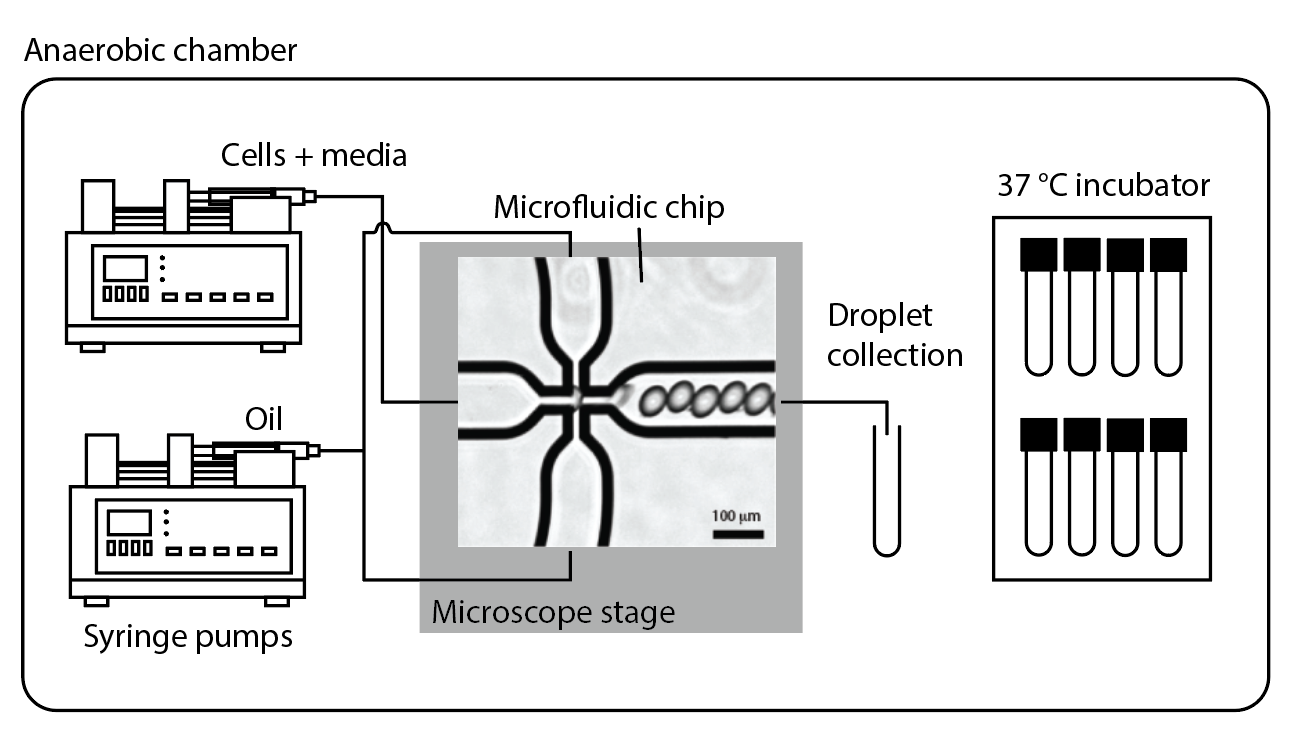

**Supplemental Figure 2.** **Schematic of droplet production in an anaerobic chamber**. Cells are encapsulated in droplets, which are formed by flowing the aqueous bacterial suspension through an immiscible oil via a T-junction on a microfluidic chip (center). Flow is controlled by two syringe pumps (left). Droplet production may be monitored by a microscope equipped with an LCD display (center). After droplets are generated, they are incubated anaerobically (right) until destructive sampling.

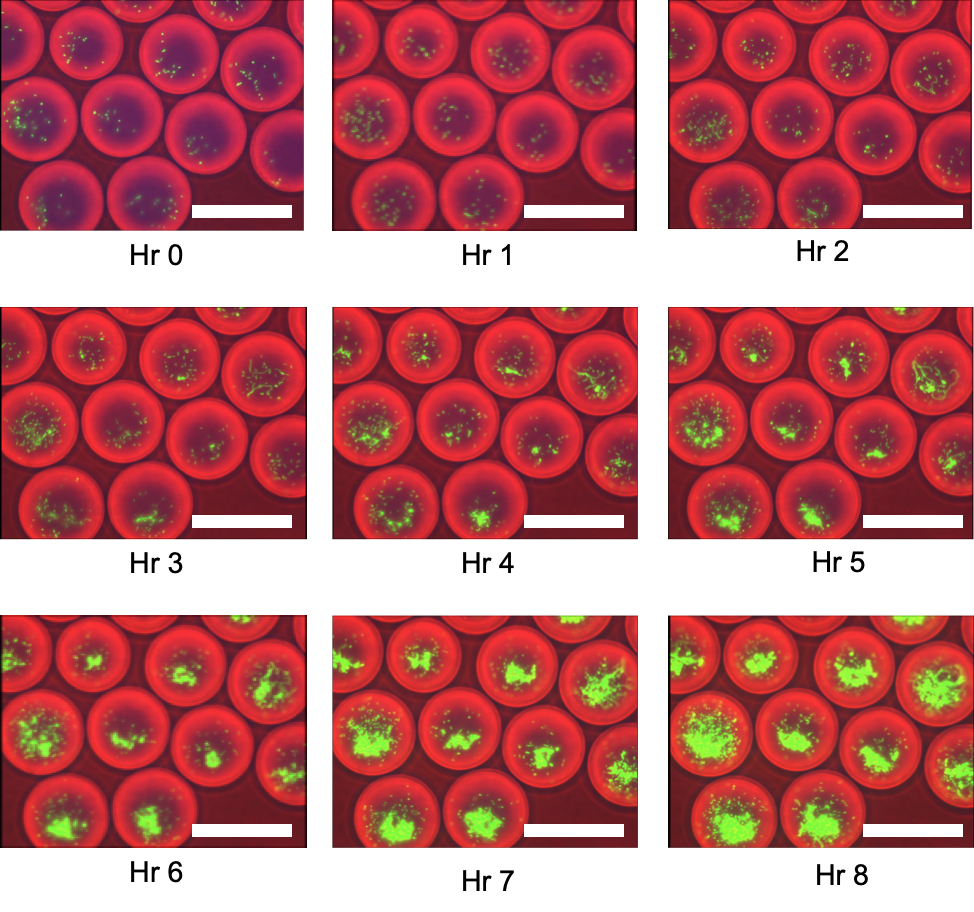

**Supplemental Figure 3.** Fluorescently labeled *E. coli* growing in droplets from 0-8 hours. In these MicDrop experiments we facilitated imaging by loading *E. coli* at high concentrations (*i.e.* most droplets were therefore loaded with more than one *E. coli* cell). Scale bars are 100 μm.

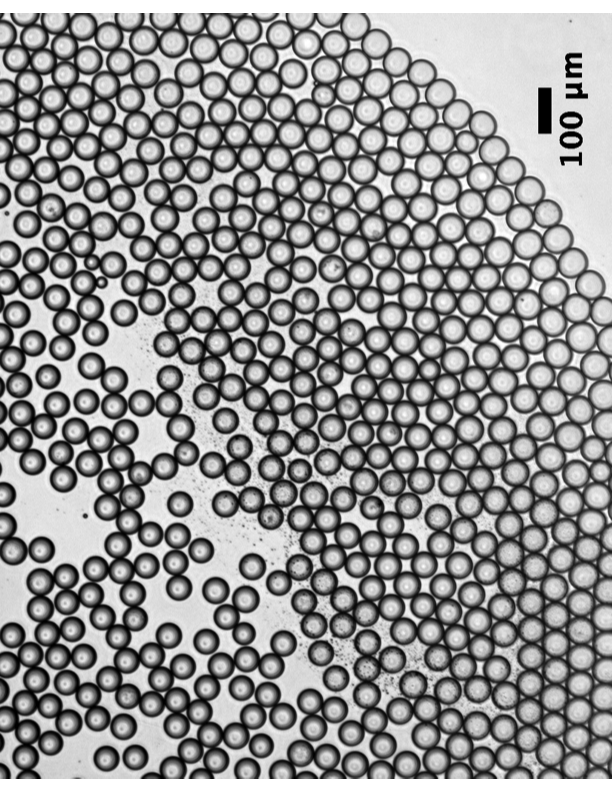

**Supplemental Figure 4.** Microfluidic droplets maintain stability and do not exhibit evidence of coalescing for at least five days. Representative image of microfluidic droplets at hour 127 (5.3 days).

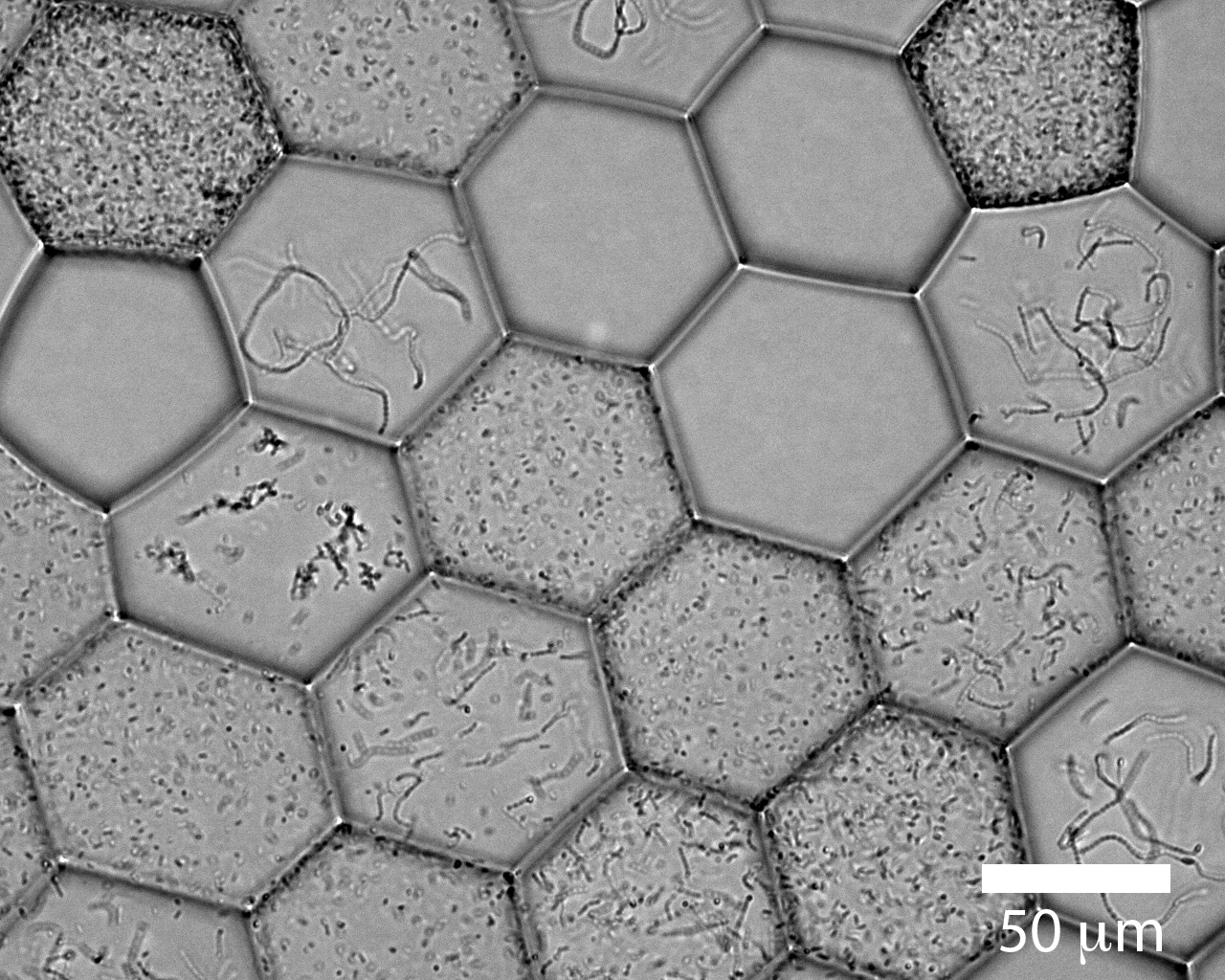
**Supplemental Figure 5. Distinct populations of bacteria observed in droplets after 6 hours incubation at 37 ℃.** An artificial community was assembled using five facultative gut anaerobes: *Streptococcus agalactiae*, *Stapholycoccus haemolyticus*, *Enterococcus faecalis*, *Enterbacter clocae*, and *Eschericia coli*. Individual isolates were: 1) grown aerobically overnight (~18 hours); 2) culture densities were determined by OD600; 3) cultures were diluted and mixed at equivalent concentrations; and, 4) mixtures were loaded into droplets. For ease of microscopic imaging, we loaded at a Poisson dilution such that > 50% of droplets contained clonal populations. To image, 5 µl of droplets were placed on glass slide, and when exposed to air the oil phase began to evaporate, and droplets became flattened hexagons on the slide.

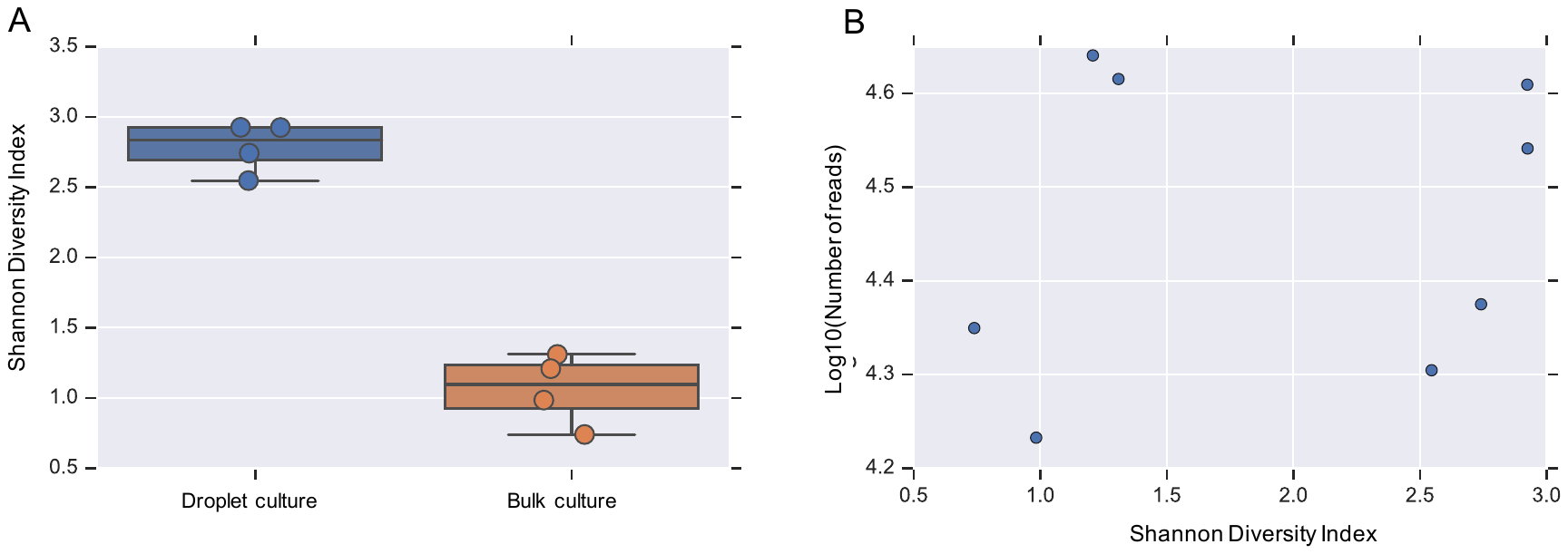

**Supplemental Figure 6.** A) Diversity (Shannon index) of microbial communities isolated and cultured in droplets culture compared with communities cultured without separation in standard bulk culture. B) Diversity (Shannon index) of these communities is not correlated with sequencing depth (y-axis, log10(number of reads)) (*p* > 0.5, Spearman correlation).

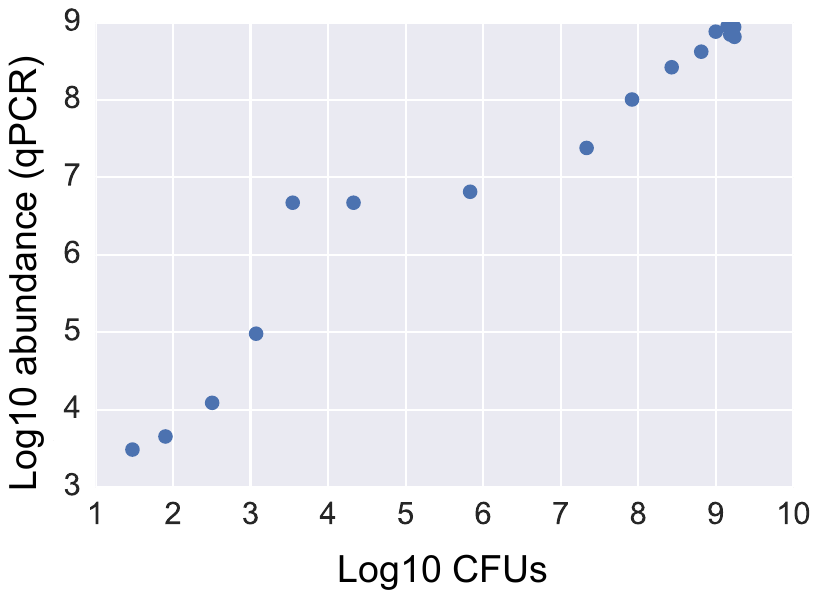

**Supplemental Figure 7.** Comparison between *E. coli* grown in plates (measured in CFUs, x-axis) and in liquid culture (measured by qPCR, y-axis; Spearman ρ=0.95, p=8.7e-9). Cultures of E. coli were grown overnight, then diluted to varying concentrations. These cultures were then simultaneously plated for CFU counting, and DNA was extracted from them for determining cell number via qPCR. These numbers were then compared, and determined that qPCR is comparable method to plate counting as a way to enumerate growing cells.

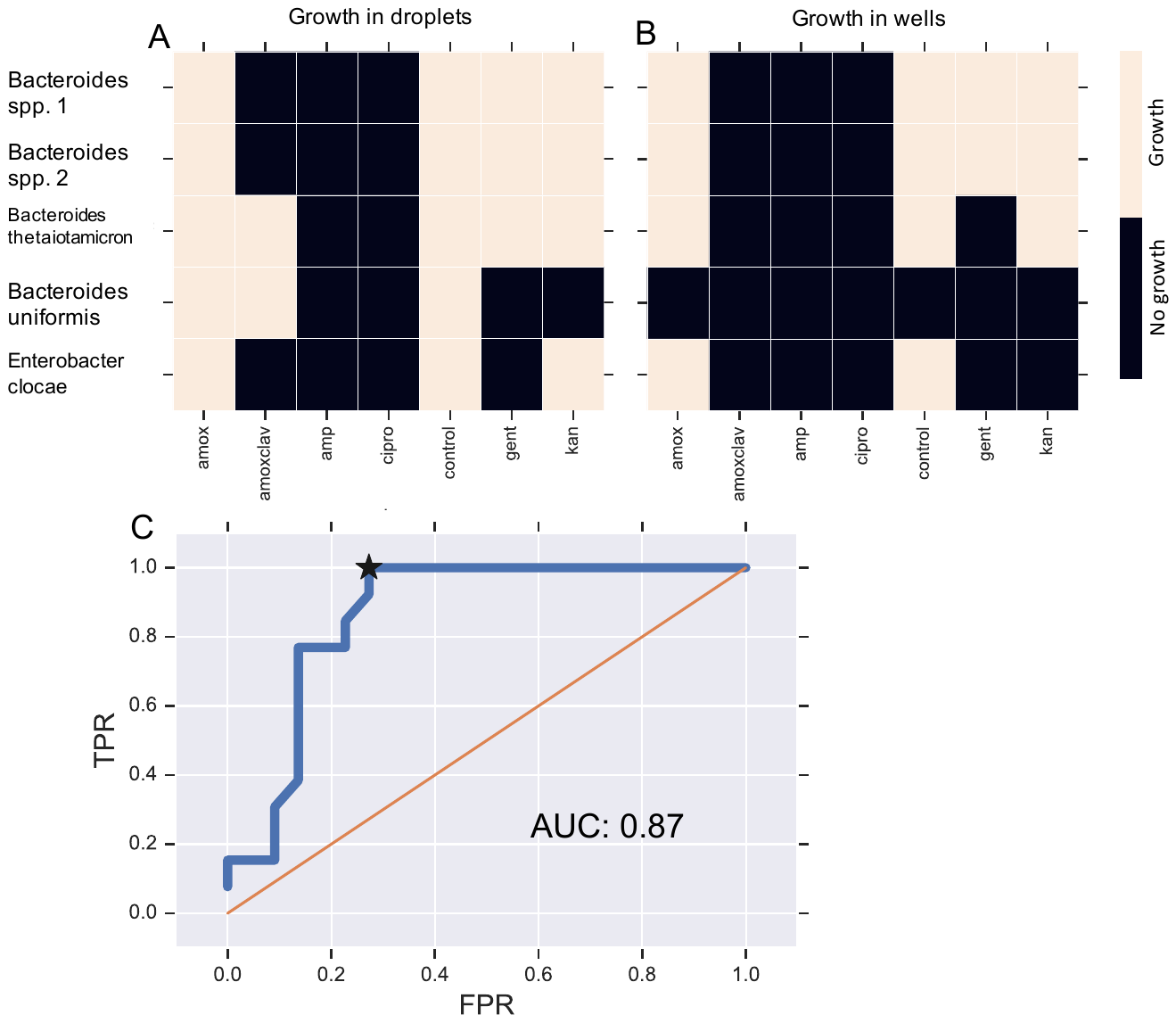

**Supplemental Figure 8.** Results of growth with antibiotics in droplets (A) and 96-well plates (B), using five gut isolates (*Bacteroides spp. 1,* *Bacteroides spp*. 2, *Bacteroides thetaiotamicron*, *Bacteroides uniformis*, *Enterobacter clocae*) grown in mGAM and six different antibiotic combinations (amoxicillin (100 µg/mL) [amox], amoxicillin + clavulanate (100 µg/mL) [amoxclav], ampicillin (100 µg/mL) [amp], gentamicin (10 µg/mL) [gent], kanamycin (50 µg/mL) [kan], and ciprofloxacin (5 µg/mL) [cipro]). Growth was measured via qPCR and sequencing (droplets) and OD600 (96-well plates) after 24 hours. (C) ROC curve of MicDrop assay results at different growth threshold cut-offs using (B) as a reference. A growth cut-off of doubling at least 2.14 times (∆ ln(SV DNA abundance) ≥ 1.48) maximized the true positive rate while minimizing the false positive rate (Youden’s J; denoted by star on curve) and was used to draw the heatmap in (A).

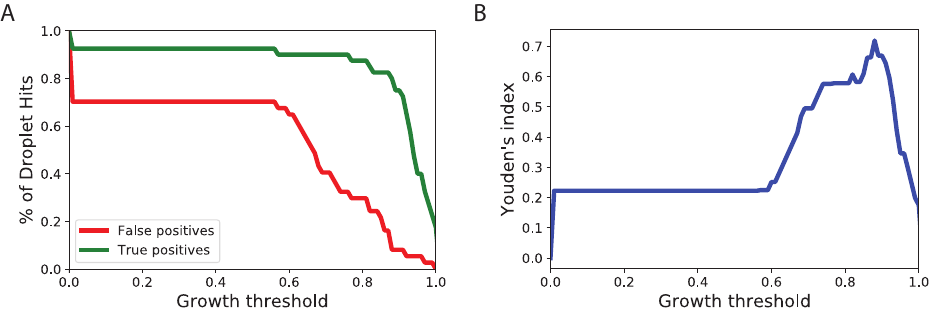

**Supplemental Figure 9.** Threshold determination for droplet prebiotic screen using an artificial community grown both in droplets and well-plates. (A) False positive and true positive rates as a function of SV growth threshold in droplets (where growth is normalized to the maximum growth value for each SV). (B) A threshold of 0.88 maximized Youden’s J index, which equally weighs both assay sensitivity and specificity, also referred to as the true positive and true negative rate, respectively.

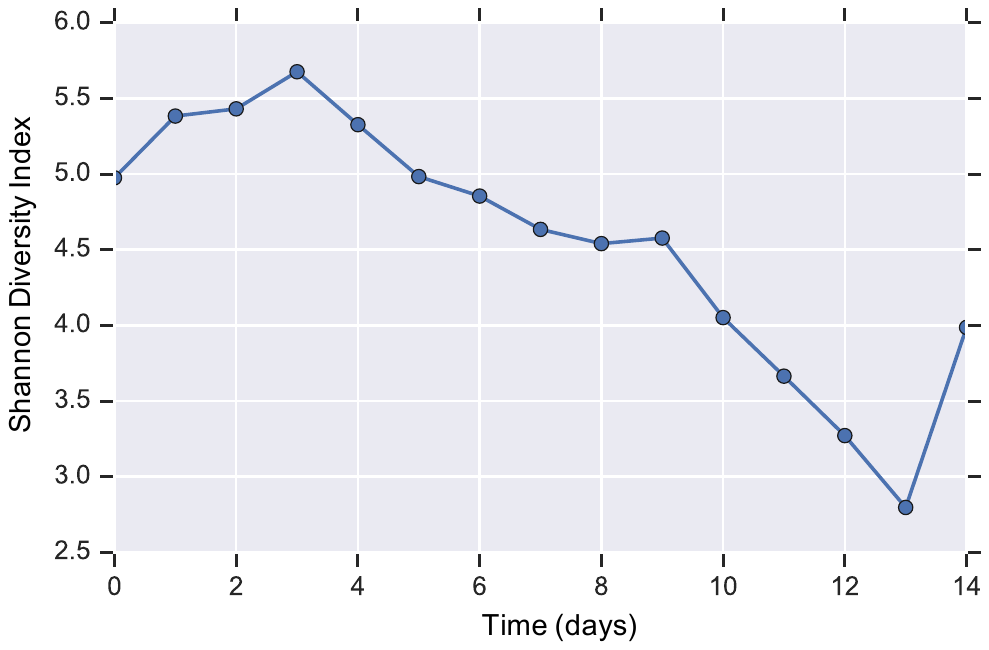

**Supplemental Figure 10.** Microbiota diversity (Shannon index) over time in the artificial gut system. (ρ = -0.91, *p* = 3.06e-6, Spearman correlation)

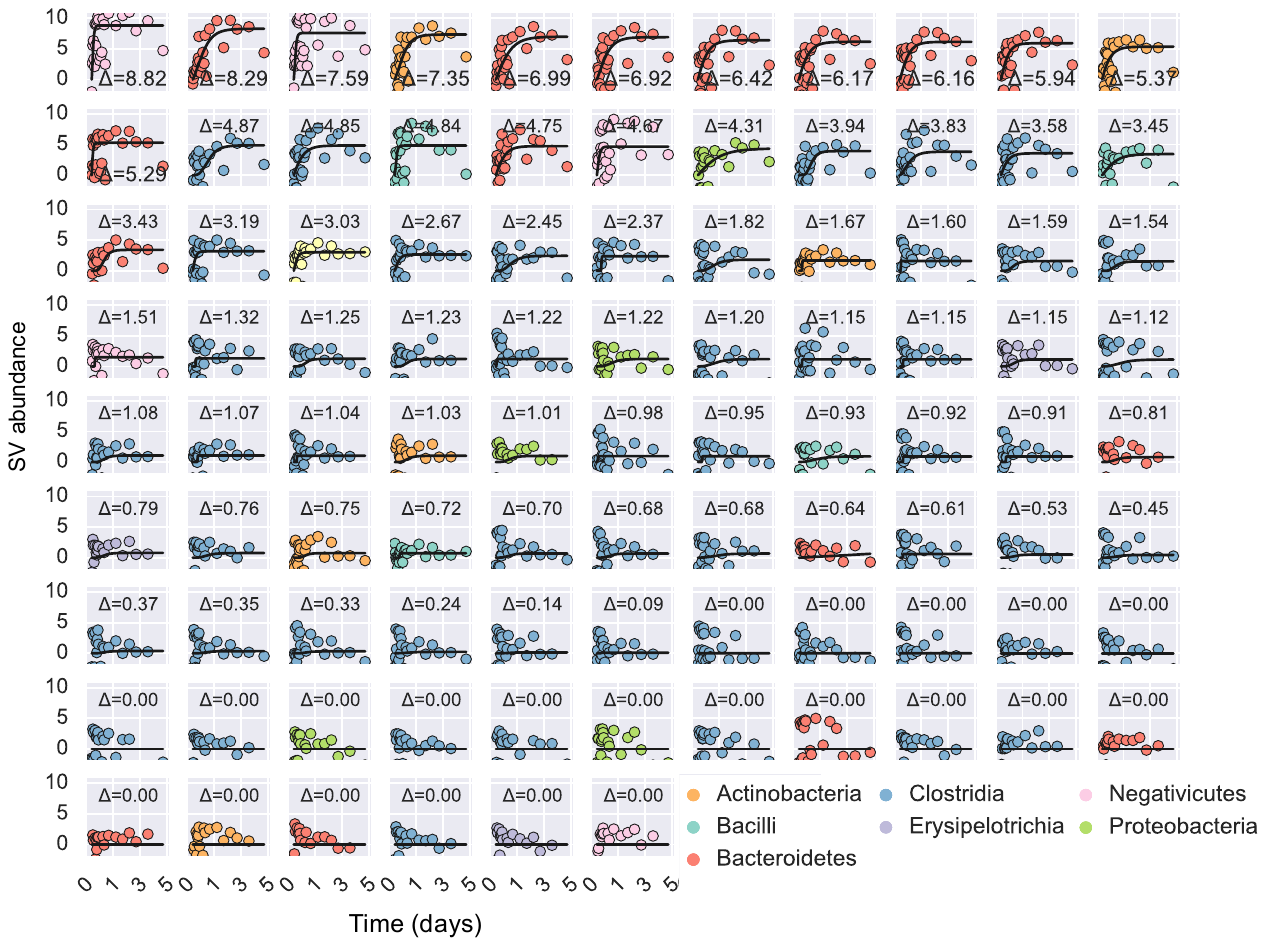

**Supplemental Figure 11.** All detected SVs in MicDrop experiment using a human stool sample. Modified Gompertz growth curves are fit to time-series. SVs are colored by taxonomy and sorted according to total growth (curve asymptote height; indicated by ∆), which is denoted on each sub-plot. To ease viewing, curves are shifted vertically so y-intercepts are at the origin.

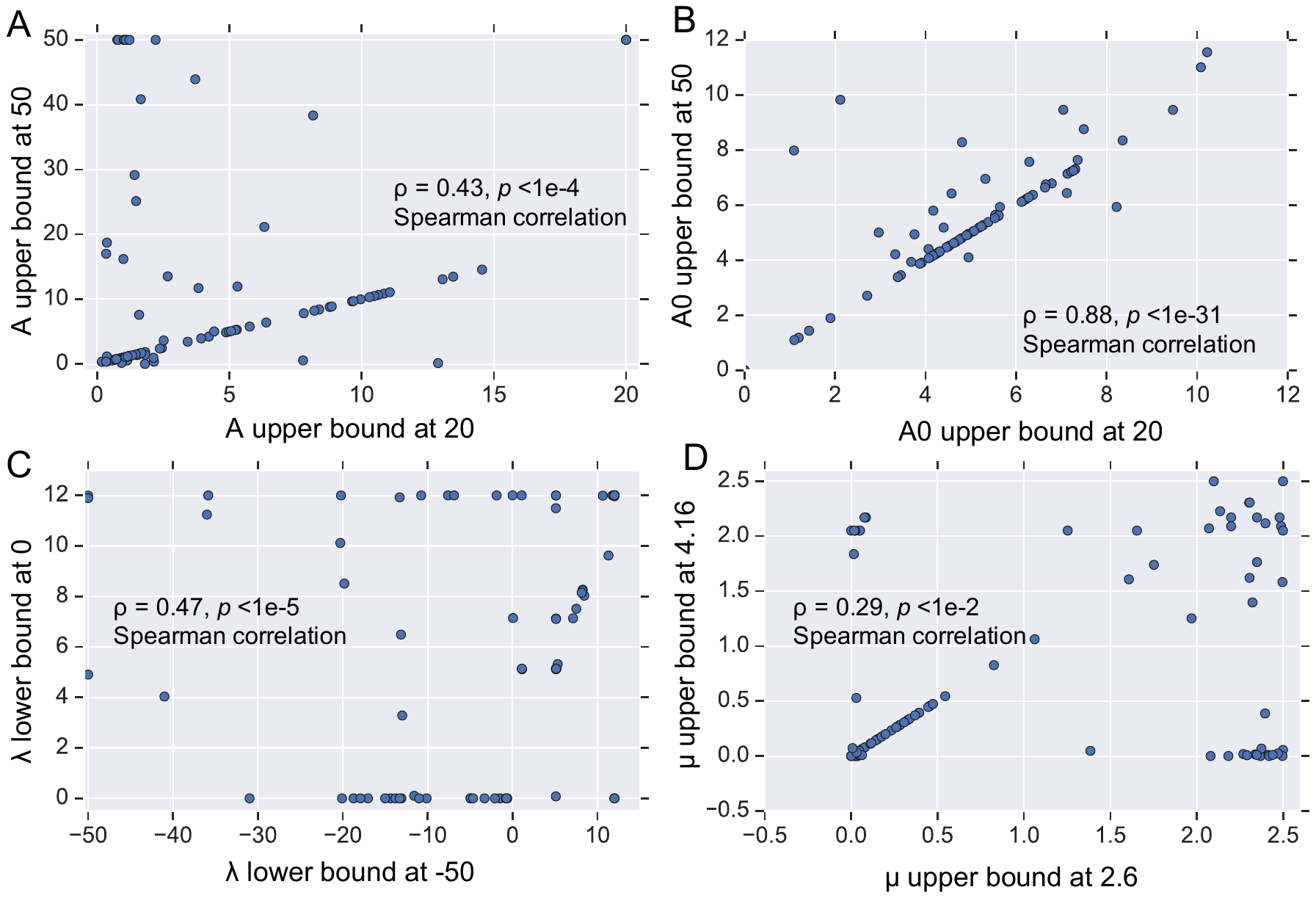

**Supplemental Figure 12.** Sensitivity of curve fitting to parameter bounds. X-axis depicts each fit parameter as defined by bounds described in *Methods*. Y-axis reflects fit parameters with alternative parameter bounds. Each point represents a fit parameters for distinct SV.

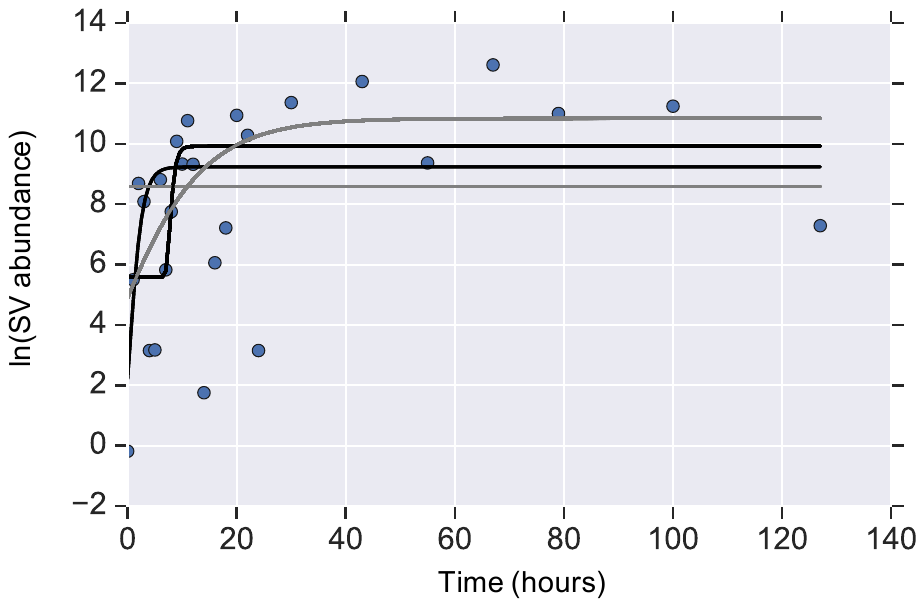

**Supplemental Figure 13. Example of optimal curve fitting**. Gompertz curves were fit to longitudinal SV data using starting parameters that were randomly chosen from bounded uniform distributions. Fitted curves typically converged on a small number of distinct local minima (solid lines). We discarded fitted curves whose growth rates (µ) collapsed to our upper bound on µ (black lines).

### Supplemental Table 1. Artificial community binary classification performance metrics. MCC denotes the Matthews correlation coefficient, which ranges from -1 to 1, and 1 being a perfect predictor.

| **Sample** | **Accuracy** | **Sensitivity** | **Specificity** | **Precision** | **FPR** | **FNR** | **FDR** | **MCC** |
| --- | --- | --- | --- | --- | --- | --- | --- | --- |
| Artificial Community | 0.869 | 0.800 | 0.931 | 0.914 | 0.068 | 0.200 | 0.085 | 0.741 |

**Supplemental Table 2.** Primary degrader richness results table by participant. MAD denotes the median absolute deviation.

| **Participant** | **Median** | **MAD** | **Min.** | **Max.** |
| --- | --- | --- | --- | --- |
| D | 23.0 | 5.5 | 11.0 | 31.0 |
| A | 21.5 | 6.0 | 15.0 | 34.0 |
| H | 13.5 | 5.0 | 8.0 | 25.0 |
| G | 14.0 | 2.8 | 11.0 | 20.0 |
| B | 12.5 | 3.3 | 6.0 | 16.0 |
| F | 11.5 | 4.8 | 5.0 | 21.0 |
| I | 7.0 | 3.3 | 5.0 | 15.0 |
| C | 7.0 | 6.8 | 4.0 | 24.0 |
| E | 5.5 | 4.0 | 4.0 | 15.0 |

**Supplemental Table 3.** PERMANOVA of primary degraders by subject and prebiotic.

|  | **DF** | **Model F** | **R^2^** | **p** |
| --- | --- | --- | --- | --- |
| Subject | 8.00 | 6.68 | 0.30 | 0.001 |
| Prebiotic | 3.00 | 9.48 | 0.16 | 0.001 |

**Supplemental Table 4.** **Fraction of inoculating stool microbes present in artificial gut over time**.

|  | TAXA # in FECAL INOCULUM | FRACTION IN ARTIFICIAL GUT | | | |
| --- | --- | --- | --- | --- | --- |
|  |  | **Day 0** | **Day 1** | **Day 7** | **Day 14** |
| Phylum | 4 | 1.00 | 1.00 | 1.00 | 1.00 |
| Class | 10 | 1.00 | 0.90 | 0.90 | 0.80 |
| Order | 10 | 1.00 | 0.90 | 0.90 | 0.80 |
| Family | 16 | 0.94 | 0.88 | 0.81 | 0.63 |
| Genus | 59 | 0.92 | 0.88 | 0.40 | 0.23 |
| Sequence Variants | 112 | 0.79 | 0.78 | 0.29 | 0.18 |

**Supplemental Table 5.** Sequence variants and taxonomy of bacteria detected in MicDrop growth experiments but not detected in inoculating stool.

|  | Phylum | Class | Order | Family | Genus | Species |
| --- | --- | --- | --- | --- | --- | --- |
| seq_1 | Bacteroidetes | Bacteroidia | Bacteroidales | Bacteroidaceae | Bacteroides |  |
| seq_107 | Firmicutes | Negativicutes | Selenomonadales | Veillonellaceae | Veillonella |  |
| seq_131 | Actinobacteria | Actinobacteria | Bifidobacteriales | Bifidobacteriaceae | Bifidobacterium | bifidum |
| seq_136 | Firmicutes | Negativicutes | Selenomonadales | Veillonellaceae | Veillonella |  |
| seq_220 | Firmicutes | Bacilli | Lactobacillales | Streptococcaceae | Streptococcus |  |
| seq_254 | Firmicutes | Clostridia | Clostridiales | Ruminococcaceae |  |  |
| seq_311 | Cyanobacteria | Chloroplast |  |  |  |  |
| seq_345 | Firmicutes | Clostridia | Clostridiales | Ruminococcaceae |  |  |
| seq_394 | Actinobacteria | Actinobacteria | Corynebacteriales | Corynebacteriaceae |  |  |
| seq_46 | Firmicutes | Negativicutes | Selenomonadales | Veillonellaceae | Veillonella |  |
| seq_48 | Bacteroidetes | Bacteroidia | Bacteroidales | Porphyromonadaceae | Parabacteroides | distasonis |
| seq_8 | Firmicutes | Bacilli | Lactobacillales | Enterococcaceae | Enterococcus |  |

**Supplemental Table 6. Minimal Medium Formulation**

Recipe to make 2X Minimal Medium to be added to 2X carbon stock, adapted from^3^ and^4^. Pre-made vitamin, trace elements and amino acid solutions were incorporated to enhance reproducibility.

| **Component** | **2X Concentration (M)** | **Source** | **Recipe** |
| --- | --- | --- | --- |
| Monopotassium phosphate | 2.0e-1 | Sigma | Martens *et al.* 2008 |
| Sodium chloride | 3.0e-2 | Sigma | Martens *et al.* 2008 |
| Ammonium sulfate | 1.7e-2 | Sigma | Martens *et al.* 2008 |
| L-cysteine | 8.0e-3 | Sigma | Martens *et al.* 2008 |
| Hematin | 3.8e-6 | Sigma | Martens *et al.* 2008 |
| L-histidine | 4.0e-4 | Sigma | Martens *et al.* 2008 |
| Magnesium chloride | 2.0e-4 | Sigma | Martens *et al.* 2008 |
| Ferrous sulfate | 2.8e-6 | Sigma | Martens *et al.* 2008 |
| Calcium chloride | 1.0e-4 | Sigma | Martens *et al.* 2008 |
| Menadione | 1.2e-5 | Sigma | Martens *et al.* 2008 |
| Cobalamin | 8.0e-9 | Sigma | Martens *et al.* 2008 |
| Vitamins and Minerals | 2X working concentration | ATCC | Adapted from Hehemann *et al*. 2010 |
| Trace Elements | 2X working concentration | ATCC | Adapted from Hehemann *et al.* 2010 |
| Amino Acid Supplement | 2X working concentration | Sigma AA-5550 | Adapted from Hehemann *et al.* 2010 |
| Purine and Pyramidine solution | 2X working concentration | Sigma | Hehemann *et al.* 2010 |
| Sodium hydroxide to pH 7.0 | - |  |  |

**Supplemental Table 7**. List of strains used in validation experiments.

| **Species** | **Usage** | **Source** |
| --- | --- | --- |
| *Bacteroides thetaiotaomicron ATCC 29148* | Prebiotic monoculture validation | ATCC |
| *Bacteroides ovatus* | Prebiotic synthetic community | Fecal isolate |
| *Ruminococcus gnavus* | Prebiotic synthetic community | Fecal isolate |
| *Escherichia coli* | Prebiotic synthetic community | Fecal isolate |
| *Klebsiella granulomatis* | Prebiotic synthetic community | Fecal isolate |
| *Bacteroides vulgatus* | Prebiotic synthetic community | Fecal isolate |
| *Enterococcus faecalis* | Prebiotic synthetic community | Fecal isolate |
| *Bacteroides 53121* | Prebiotic synthetic community | Fecal isolate |
| *NDL-177, Eschericia coli* | Fluorescent droplet experiments | Gift of N. Lord |
| *Streptococcus agalacticaeae* | Validation synthetic community | Fecal Isolate |
| *Bacteroides fragilis* | Validation synthetic community | Fecal Isolate |
| *Bifidobacterium longum* | Validation synthetic community | Fecal Isolate |

**Supplemental Table 8.** Carbon sources used in this work.

| **Carbon source** | **Cat. No.** | **Source** | **Experiment** |
| --- | --- | --- | --- |
| Pullulan | P4516 | Sigma | Monoculture validation + Artificial community |
| Levan | L8647 | Sigma | Monoculture validation |
| Lamanarin | L0088 | TCI | Monoculture validation + Artificial community |
| Arabinogalactan | A1333 | Spectrum | Artificial community |
| Arabinose | A3256 | Sigma | Artificial community |
| FOS | F8052 | Sigma | Artificial community |
| Fructose | 0226 | Amresco | Artificial community |
| Galactose | G0750 | Sigma | Artificial community |
| Glucose | 0188 | Amresco | Artificial community |
| Inulin | I3754 | Sigma | Artificial community |
| Mannose | M6020 | Sigma | Artificial community |
| Xylan | P-XLYNBE | Megazyme | Human donors + Artificial Community |
| GOS | Bimuno Daily | Clasado Biosciences | Human donors |
| Inulin | F97 | Cargill | Human donors |
| Dextrin | Benefiber | GSK | Human donors |
